# Angiogenin activity regulates RNA remodeling during the maternal-to-zygotic transition in early mouse embryogenesis

**DOI:** 10.64898/2026.09.22.753398

**Authors:** Shikha Sharma, Nivedita Hariharan, Ambika S. Kurbet, Praveen Kumar Vemula, Aurélie Jory, Tina Mukherjee, Dasaradhi Palakodeti

## Abstract

Angiogenin (ANG), a member of the RNase A superfamily, regulates multiple aspects of RNA metabolism, including rRNA processing and the generation of tRNA-derived small RNAs. However, its function during early mammalian embryogenesis remains unknown. Here, we identify a previously unrecognized requirement for ANG activity during early mouse embryonic development. Pharmacological inhibition of ANG during the first embryonic cell cycle did not prevent progression to the 2-cell stage but markedly impaired subsequent development to the 4-cell stage, indicating a critical requirement for ANG activity during the maternal-to-zygotic transition. Transcriptome-wide analysis revealed that ANG inhibition disrupted the normal remodeling of the embryonic transcriptome, characterized by persistent accumulation of a subset of transcripts that are normally reduced during the 2-cell transition, including many maternally deposited RNAs, together with altered expression of transcripts associated with zygotic genome activation. Small-RNA profiling further revealed widespread alterations in miRNA and tRNA-derived small-RNA populations. Notably, a large proportion of transcripts retained following ANG inhibition contained predicted target sites for miRNAs that were concomitantly reduced, suggesting a potential link between ANG-dependent small-RNA regulation and maternal transcript clearance. ANG inhibition was also associated with altered abundance and organization of components of RNA regulatory granules, including Tudor, TIAR, Dcp1a and Ago2, and increased association of selected retained transcripts Reep4 and Esrrb with Tudor-positive structures. Together, these findings identify ANG activity as an important regulator of RNA remodeling during the maternal-to-zygotic transition and suggest that ANG coordinates maternal transcript clearance, small-RNA dynamics and mRNP organization to support early mouse embryonic development.

## Introduction

Mature mammalian oocytes are transcriptionally quiescent and therefore depend on maternally deposited RNAs and proteins to support fertilization and the earliest stages of embryonic development [1,2]. Following fertilization, developmental control is progressively transferred from maternal gene products to the newly activated embryonic genome through the maternal-to-zygotic transition (MZT). In mice, zygotic transcription begins during the mid-to-late 1-cell stage as minor zygotic genome activation (minor ZGA) and is followed by a major wave of transcription at the 2-cell stage, referred to as major ZGA [3]. Successful MZT requires coordinated activation of the zygotic genome together with extensive remodeling of the maternally inherited transcriptome. Maternal transcripts are selectively cleared in a temporally regulated manner through both maternally encoded decay mechanisms and pathways coupled to zygotic transcription [1,2]. Disruption of either maternal RNA clearance or ZGA can compromise developmental progression, highlighting the importance of coordinated RNA turnover during early embryogenesis. However, the full repertoire of RNA-processing factors that regulate this transition remains incompletely understood.

Angiogenin (ANG) is a secreted ribonuclease belonging to the RNase A superfamily that has been implicated in diverse biological processes, including angiogenesis, cellular proliferation, stress responses, rRNA transcription and RNA metabolism [4–8]. Although structurally related to RNase A, ANG displays substantially lower ribonucleolytic activity and distinct substrate preferences, suggesting that its biological functions depend on regulated recognition and processing of specific RNA substrates [9–15]. A particularly well-characterized function of ANG is its ability to cleave tRNAs under cellular stress, generating tRNA-derived stress-induced RNAs (tiRNAs) that can modulate translation and stress responses [4,16–19]. ANG has also been implicated in transcriptional regulation and other aspects of RNA metabolism [7], suggesting that its functions extend beyond tRNA cleavage. Thus, ANG is positioned at the interface between RNA processing, small-RNA generation and cellular adaptation.

The extensive remodeling of the RNA landscape that accompanies the MZT raises the possibility that regulated ribonuclease activity may contribute to early embryonic development. However, whether ANG participates in this process has not been investigated. Here, we examined the requirement for ANG activity during early mouse embryogenesis using pharmacological inhibition of its ribonucleolytic activity. We find that inhibition of ANG during the first embryonic cell cycle permits development to the 2-cell stage but markedly compromises subsequent progression to the 4-cell stage, indicating an early requirement for ANG activity. Transcriptome-wide analyses reveal that ANG inhibition disrupts the normal remodelling of the embryonic transcriptome, including the persistence of a subset of maternally deposited transcripts and altered expression of transcripts associated with ZGA. Small-RNA profiling further identifies changes in miRNA and tRNA-derived small-RNA populations, while analysis of RNA regulatory granule components reveals altered abundance and organization of factors associated with mRNA turnover. Together, these findings identify a previously unrecognized requirement for ANG activity during the maternal-to-zygotic transition and suggest that ANG contributes to the coordinated RNA remodeling required for early mouse embryonic development.

## Results

### Maternal transcript dynamics during early mouse embryogenesis

To characterize the dynamics of maternally deposited transcripts during early mouse embryogenesis, we compared transcriptomic profiles of unfertilized oocytes (UFO), fertilized oocytes (FO), late 1-cell (L-1c), 2-cell (2c), and 4-cell (4c) stage embryos (Figure 1A, Supp Figure 1A, Supp Table S1). Transcripts that were abundant in UFO and FO and progressively decreased during development from the 1-cell to the 4-cell stage were classified as maternal transcripts, consistent with previously described patterns of maternal RNA clearance during the maternal-to-zygotic transition [1,20]. We further shortlisted transcripts that were enriched by more than two-fold in UFO relative to the 1-cell, 2-cell, and 4-cell stages (Figure 1B). Analysis of their temporal expression profiles revealed a progressive reduction in the abundance of these maternal transcripts following fertilization, with a pronounced decline during the late 1-cell and subsequent 2-cell and 4-cell stages (Figure 1B, Supp Figure 1A, Supp Table S1).

**Figure 1:**
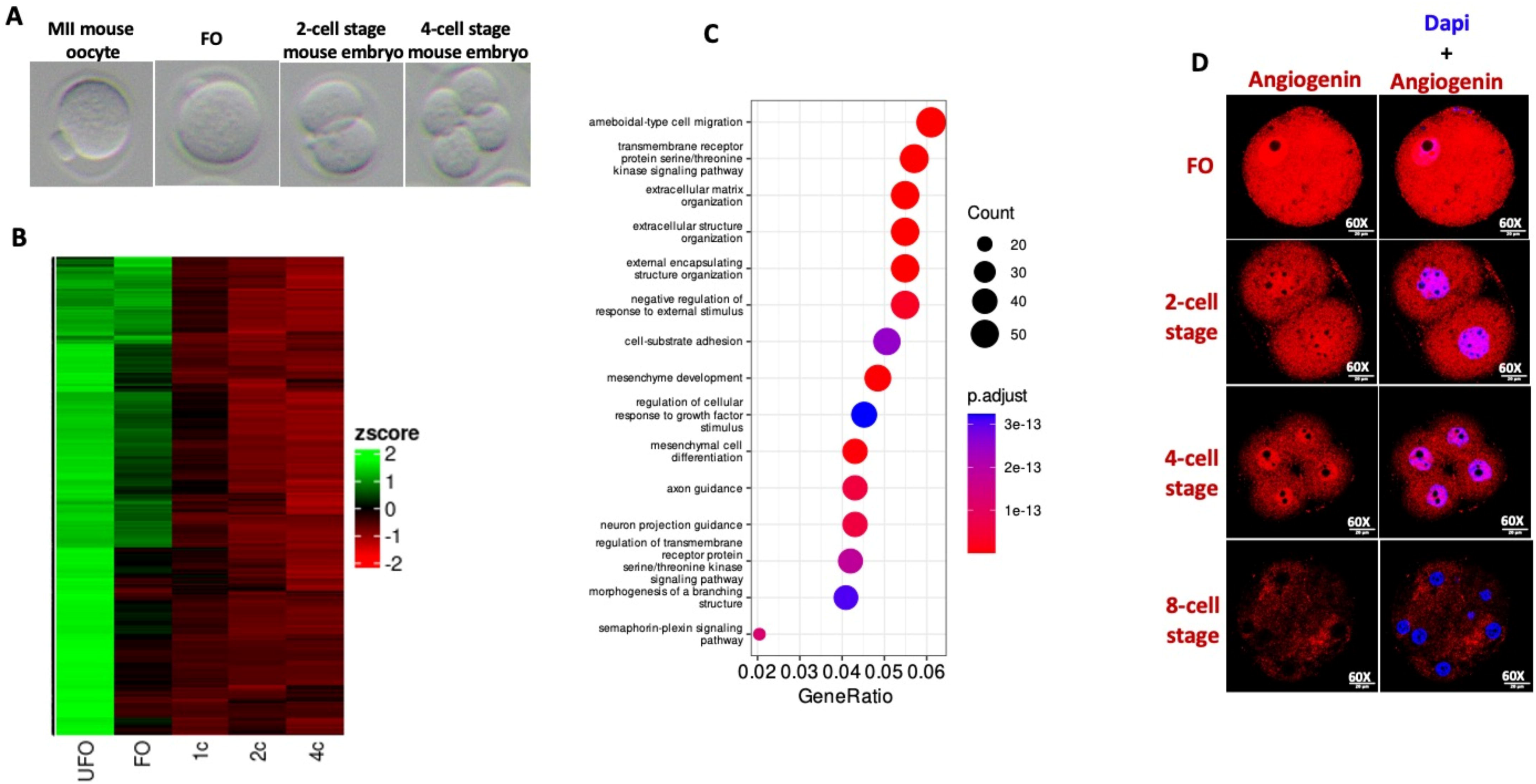
**A)** DIC image representation of MII oocytes, FO, 2-cell stage and 4-cell stage mouse embryos B) Heatmap depicts the maternal RNA degradation pattern during early developmental stages of m-emb at FO, 1c, 2c and 4c compared to UFO. C) GOBP analysis of maternal RNAs represented in heat map B D) Representative images shows immunofluorescence detection of angiogenin expression in FO, 2-cell stage, 4-cell stage of m-embs. **Note: UFO = unfertilized oocytes, FO = fertilized oocytes, L-1c = late 1-cell, 2c = 2-cell stage, and 4c = 4-cell stage**

Gene Ontology analysis of the maternal transcripts represented in the heatmap identified enrichment of biological processes and molecular functions associated with serine/threonine kinase activity, extracellular structure organization, and mesenchymal and neuronal developmental programs (Figure 1C). These findings illustrate the extensive remodeling of the maternally inherited transcriptome that accompanies early embryonic development.

We next examined ANG expression during the same developmental window. ANG expression was highest in fertilized oocytes and declined during subsequent development, with lower expression observed at the 2-cell, 4-cell, and 8-cell stages (Figure 1D, Supp Figure 1B). Thus, ANG is prominently expressed during the developmental period in which extensive remodelling of the maternal transcriptome occurs. This temporal association prompted us to investigate whether ANG activity contributes to maternal transcript clearance and early embryonic progression.

### Angiogenin activity is required for efficient progression of mouse embryos from the 2-cell to the 4-cell stage

To investigate the requirement for ANG activity during early mouse embryogenesis, fertilized oocytes (FO) were treated with 50 μM angiogenin inhibitor (ANGi). Under our culture conditions, embryos progressed from the 1-cell to the 2-cell stage within approximately 18–20 h and reached the 4-cell stage by 47 ± 1 h after fertilization. Since ANGi was dissolved in DMSO, embryos treated with an equivalent concentration of DMSO were used as controls. While DMSO-treated embryos progressed to the 4-cell stage, continuous exposure to ANGi resulted in a pronounced developmental arrest at the 2-cell stage at 47 ± 1 h (Figure 2A). These observations indicate that inhibition of ANG activity compromises developmental progression from the 2-cell to the 4-cell stage.

**Figure 2:**
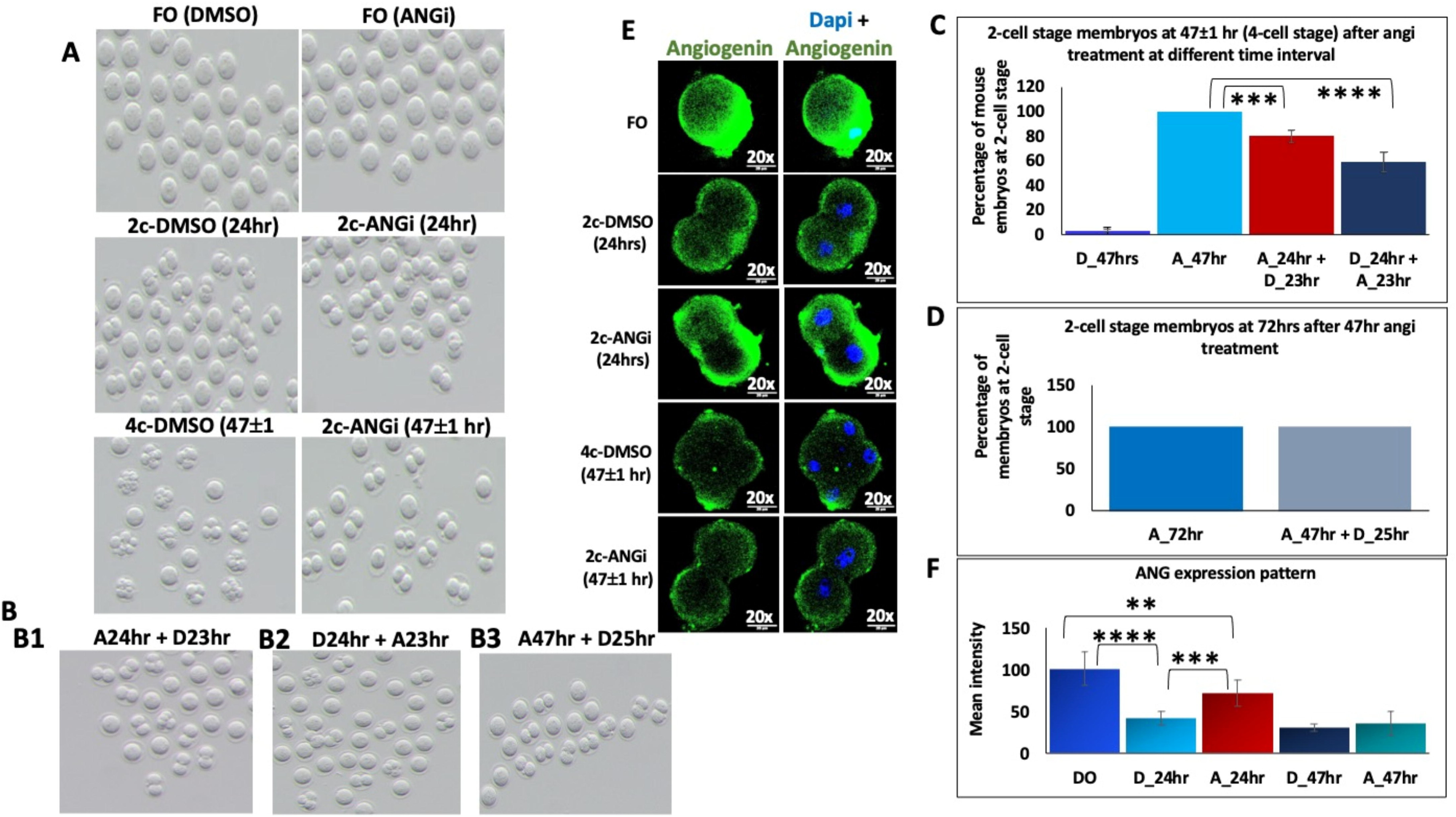
A) Representative bright-field images of m-embs cultured in the presence and absence of ANGi for 24hrs and 47±1 hrs compared to the control condition. B) B1) Representative bright-field image of m-embs at 47±1 hrs cultured in the presence of ANGi for 24hrs immediately after FO isolation and further replaced with control media for 23 hr B2) Representative bright-field image of m-embs at 47±1 hrs cultured in the presence of DMSO control for 24hrs after FO isolation and further replaced with ANGi media for 23 hr B3) Representative bright-field image of m-embs at 72 hrs cultured in the presence of ANGi for 47 hrs after FO isolation and further replaced with DMSO control media for 25 hrs. C) The Bar graph denotes the number of m-embs struck at the 2-cell stage upon ANGi treatment at different time intervals calculated from the above A and B experiments. The experiment was done n=3 times D) The Bar graph shows the number of m-embs struck at the 2-cell stage upon ANGi treatment at different time intervals calculated from the above experiment (B3). The experiment was done n=3 times. E) Representative images shows immunofluorescence detection of ANG expression in m-embs cultured in the presence and absence of ANGi for 24hrs and 47±1 hrs compared to DMSO control condition. The experiment was done n=3 times F) Bar graph represents the mean intensity of ANG expression in m-embs upon ANGi treatment for 24hrs and 47±1 hrs compared to DMSO control. The mean intensity was calculated from the confocal images (n=11-13) using ImageJ software. The experiment was done n=3 times. **Note: D=DMSO, A= ANGi** ***Statistical analysis was performed using the two-sample t-test method, with the mean and standard deviation calculated as described by Xu et al. (2017). The p-values were calculated using MedCalc online software (MedCalc Software, 2024). P-value are represented with asterisk <0.05 (*), <0.01 (**), <0.001 (***), <0.0001 (****)*.**

We next investigated whether the developmental outcome depended on the timing of ANG inhibition. Fertilized oocytes were exposed to ANGi for the first 24 h of development and subsequently cultured in inhibitor-free medium. Transient exposure to ANGi did not substantially affect progression from the 1-cell to the 2-cell stage during the initial 24 h; however, approximately 80% of these embryos subsequently failed to progress to the 4-cell stage by 47 ± 1 h (Figures 2B1 and 2C). This finding suggests that ANG activity during the first embryonic cell cycle is important for developmental events required for subsequent progression beyond the 2-cell stage.

To determine whether ANG activity is also required after embryos have reached the 2-cell stage, ANGi was added to 2-cell embryos approximately 24 h after fertilization. Under these conditions, approximately 60% of embryos failed to progress to the 4-cell stage by 47 ± 1 h (Figures 2B2 and 2C). Thus, inhibition initiated at the 2-cell stage also impaired developmental progression, although the effect was less pronounced than that observed when embryos were exposed to ANGi during the first 24 h of development. Together, these results suggest that ANG activity is required during early embryogenesis, with inhibition during the first embryonic cell cycle having a particularly pronounced effect on subsequent 2-cell-to-4-cell progression.

We next asked whether the developmental arrest associated with prolonged ANG inhibition was reversible. Following 47 h of ANGi treatment, embryos arrested at the 2-cell stage were transferred to inhibitor-free control medium and cultured for an additional 25 h. These embryos remained predominantly at the 2-cell stage (Figures 2B3 and 2D), indicating that developmental progression was not restored within the 25-h recovery period. Thus, under the conditions examined, removal of ANGi was insufficient to rescue the developmental arrest.

Finally, we examined ANG expression following inhibitor treatment. ANG expression decreased in both DMSO-treated and ANGi-treated 2-cell embryos relative to fertilized oocytes at 24 h and 47 ± 1 h (Figures 2E and 2F), consistent with the developmental decline in ANG expression observed in Figure 1. Interestingly, ANG levels were significantly higher in ANGi-treated embryos than in DMSO-treated embryos at 24 h. The biological significance of this difference remains unclear and will require further investigation.

### Influence of angiogenin inhibition on the maternal RNA degradation pattern

In order to understand the transcriptomic state of the embryos under developmental block, we performed transcriptome sequencing to analyze the differential expression of transcripts from 2c-ANGi and 2c-DMSO control compared to FO. The gene sets from these two comparisons (q<0.05 in at least one comparison) were then used to construct a scatter plot to delineate the mRNAs that are different between ANGi and the control samples (Figure 3A). Our analysis revealed 186 transcripts that remain either unchanged (defined as anything less than a two-fold change) or upregulated in 2c-ANGi but severely downregulated in 2c-DMSO compared to FO (Supp Table S2A). We also observed 51 transcripts that were either unchanged or upregulated in 2c-DMSO. But downregulated in 2c-ANGi compared to FO (Supp Table S2B). Following this, we constructed a heat map for the 186 transcripts to study their dynamic from FO to the 4-cell development stage of m-emb in both control and ANGi-treated m-embs for 24hrs and 47±1 hr (Figure 3B). We observed that 186 mRNAs were highly downregulated in 2c-dmso condition at 24hr and most of them became upregulated in 4c-dmso at 47±1 hr while in ANGi condition these transcripts were either retained or upregulated or degraded or were undergoing slow degradation in 47hr ±1 (Figure 3B, cluster 1, 2, 3, 4, 5), which suggest that degradation of these transcripts at 2-cell stage might be crucial for the transition of m-emb from 2-cell stage to 4-cell stage. In addition, we observed from the heat map in cluster 3 that there were sets of transcripts highly downregulated in the control condition at both 24hr and 47hr±1 and retained/upregulated in ANGi under the same condition, which exemplifies that the degradation of these mRNAs may be critical for later stages of early progression, beyond the 4-cell stage (Figure 3B). Furthermore, we noted that in clusters 4 and 5, there was delayed degradation of transcripts in the ANGi condition at 47±1 hr. In contrast, in cluster 4 there was upregulation of these transcripts in 4c-DMSO compared with 2c-ANGi at 47±1 hr, which indicates that ANG might also be playing a role in the regulation of transcription. Next, we compared the genes differentially regulated in 2c-ANGi relative to FO with the database based on the DBTMEE classification [21, 22] of transcripts from mouse early embryos as maternal deposited transcripts, transcripts belonging to minor and major zygotic genome activation (minor-ZGA, major-ZGA), mid-preimplantation genome activation (MGA). Our analysis revealed 40% of the transcripts upregulated in the ANGi-treated conditions are classified as maternal RNA, 18% to minor-ZGA, 17% to major-ZGA, and 15% MGA (Figure 3C, Supp Table S2A). Further, we also conducted the DBTMEE analysis for downregulated transcripts in 2c-ANGi and observed 31% to major-ZGA, and 29% to MGA, while the others belong to maternal mRNA (22%) and minor ZGA ( 6%) (Figure 3E, Supp Table S2B). Together, our data show high enrichment of maternal deposited RNA in the early embryo development under angiogenin inhibited conditions. GO annotation for biological process (GOBP) analysis for the 186 upregulated transcripts in 2c-ANGi revealed that most of the transcripts are critical for organelle assembly, vesicle-mediated transport, establishment of protein localization, macromolecule catabolic process, and intracellular transport (Figure 3D). Similar analysis for down-regulated transcripts in angiogenin-inhibited condition (2c-ANGi) showed enrichment for functions associated with RNA 3’end processing (Figure 3F), which could explain the enrichment of maternal RNA in embryos in angiogenin inhibitor-treated conditions.

**Figure 3:**
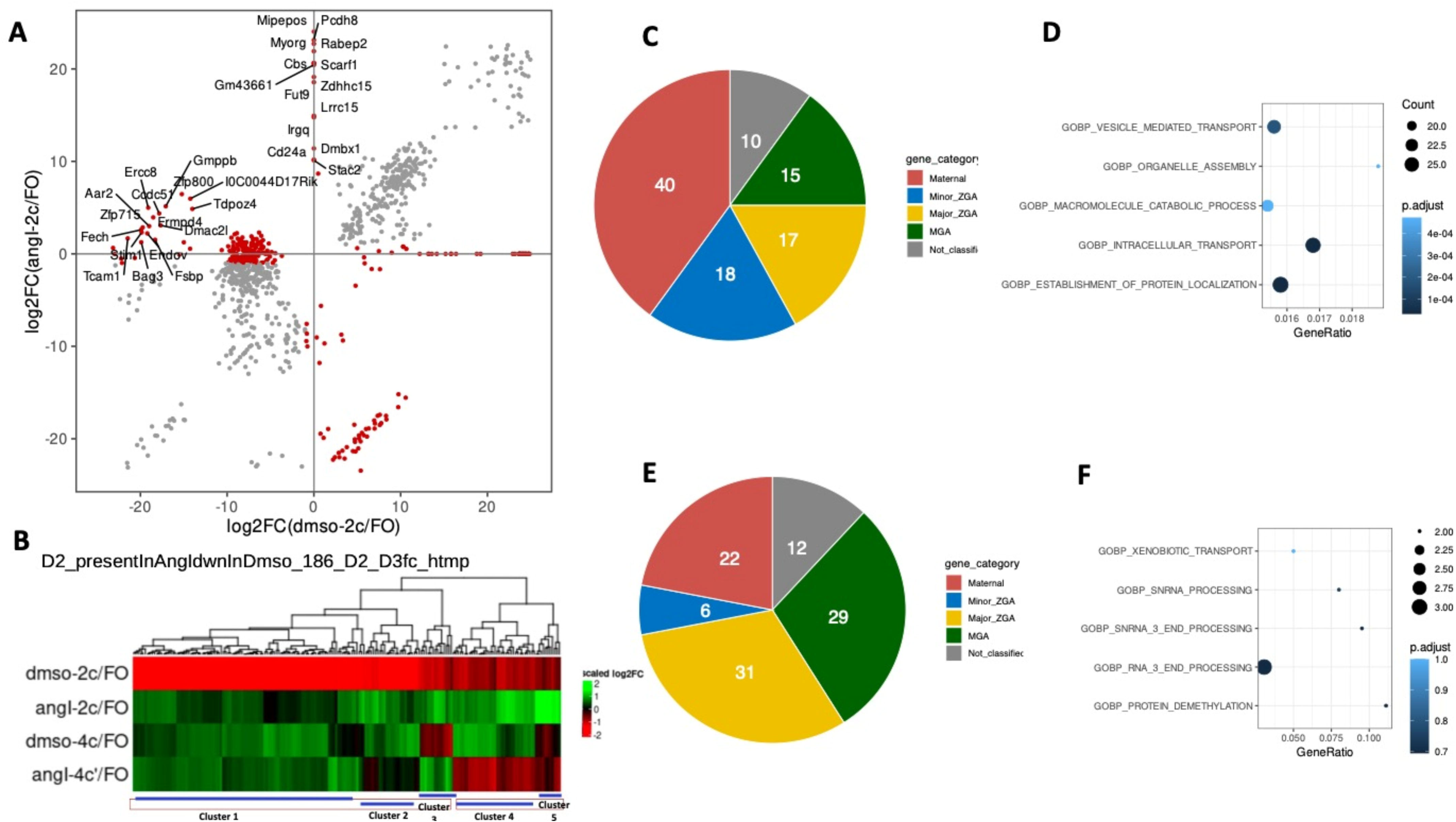
A) A Scatter plot represents the mRNAs significantly upregulated and downregulated in 2c-ANGi/FO compared to 2c-DMSO/FO condition at 24hrs. B) Heat map representation depicts the scaled log2 fold change for the 186 mRNAs that remain either unchanged or upregulated in 2c-ANGi/FO compared to 2c-DMSO/FO condition at 24hrs and further compared with 2c-ANGi/FO and 4c-DMSO/FO condition at 47±1 hrs. The heat map was plotted by combining n=2 experiments (pooled mRNA sequencing) C) DBTMEE analysis of the 186 mRNAs shown in the heat map (B). D) GOBP analysis of the 186 mRNAs shown in the heat map (B). E) DBTMEE analysis of mRNAs significantly downregulated in 2c-ANGi/FO compared to 2c-dmso/FO at 24hrs. F) GOBP analysis of mRNAs significantly downregulated in 2c-ANGi/FO compared to 2c-DMSO/FO condition at 24hrs.

To validate the RNA-seq data, which showed accumulation of mRNAs upon ANGi, we performed in situ hybridization for selected transcripts, including Reep4 (maternal RNA), Stam (minor-ZGA–MGA), Esrrb (MGA), and Map3k20 (Figure 4A, Supp Figure 2A). Compared to fertilized oocytes (FO) at 24 hours, we observed reduced expression of these transcripts in 2-cell DMSO-treated embryos (2c-DMSO), whereas expression was retained in 2-cell Angiogenin-inhibited embryos (2c-ANGi), consistent with our RNA-seq findings (Figure 4A, 4B, Supp Table S2A). Furthermore, transcripts such as Reep4, Stam, Esrrb, and Map3k20 displayed moderate to high expression levels in control embryos at 47 hours post-fertilization (47±1 h). However, in ANGi-treated embryos, these transcripts remained persistently upregulated, without any significant downregulation at 47 hours (Supp Figure 2A and 2B). This indicates a failure in the clearance of maternal RNAs, suggesting that angiogenin inhibition results in dysregulation of maternal mRNA degradation.

**Figure 4:**
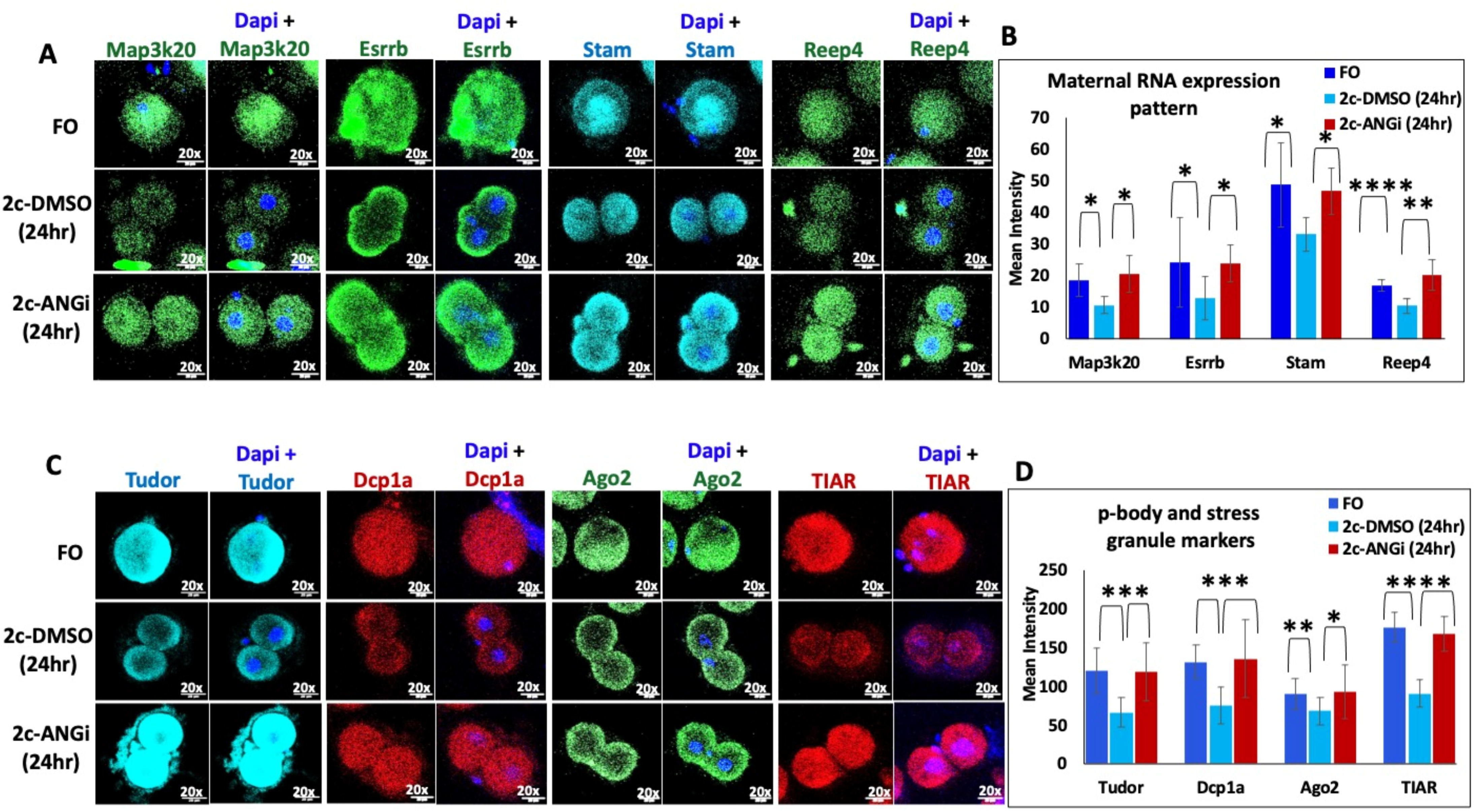
A) In-situ hybridization was done for the mRNAs detected in RNA seq data, including Map3k20, Esrrb, Stam, and Reep4 in the m-emb cultured in the presence and absence of ANGi for 24 hours, compared to the DMSO control shown by representative images. The experiments were done on the animals (n=15) obtained from the pooled samples of 3-4 breeding pair mice and approx. 15-20 mouse embryos were analysed for each condition. B) The Bar graph represents the mean intensity of Map3k20, Esrrb, Stam, and Reep4 mRNA expression in FO, 2c-DMSO, and 2c-ANGi at 24hrs The mean intensity was calculated from the confocal images (n=4-8) using ImageJ software. C) Representative images shows immunofluorescence detection of stress granules markers (Tudor, TIAR), p-body marker (Dcp1a), and RISC complex marker (Ago2) in FO, 2c-DMSO, and 2c-ANGi condition at 24hrs. The experiments were done on the animals (n=15) obtained from the pooled samples of 3-4 breeding pair mice and approx. 15-20 mouse embryos were analysed for each condition. The experiment was repeated n=3 times. D) Bar plot shows the mean intensity of Tudor, Dcp1a, Ago2, and TIAR expression in FO, 2c-DMSO, and 2c-ANGi treated m-embs at 24hrs. The mean intensity was calculated from the confocal images (n=8-16) using ImageJ software. *p-value are represented with asterisk <0.05 (*), <0.01 (**), <0.001 (***), <0.0001 (****)*.

### Influence of angiogenin inhibition on small RNA expression

To investigate whether ANG activity is associated with changes in the small RNA landscape during early embryogenesis, we performed small RNA sequencing and mapped reads ranging from 18–35 nt to the mouse genome. The proportion of reads mapping to piRNAs and miRNAs was higher in 2c-DMSO embryos at 24 h compared with unfertilized oocytes (UFO), fertilized oocytes (FO), and 2c-ANGi embryos (Figure 5A, Supp Table S3A). In contrast, the proportion of tRNA-mapped reads was lower in 2c-DMSO embryos compared with UFO, FO, and 2c-ANGi embryos at 24 h (Figure 5A, Supp Table S3A), suggesting substantial changes in the abundance of tRNA-derived small RNAs (tsRNAs) following fertilization and during progression to the 2-cell stage.

**Figure 5:**
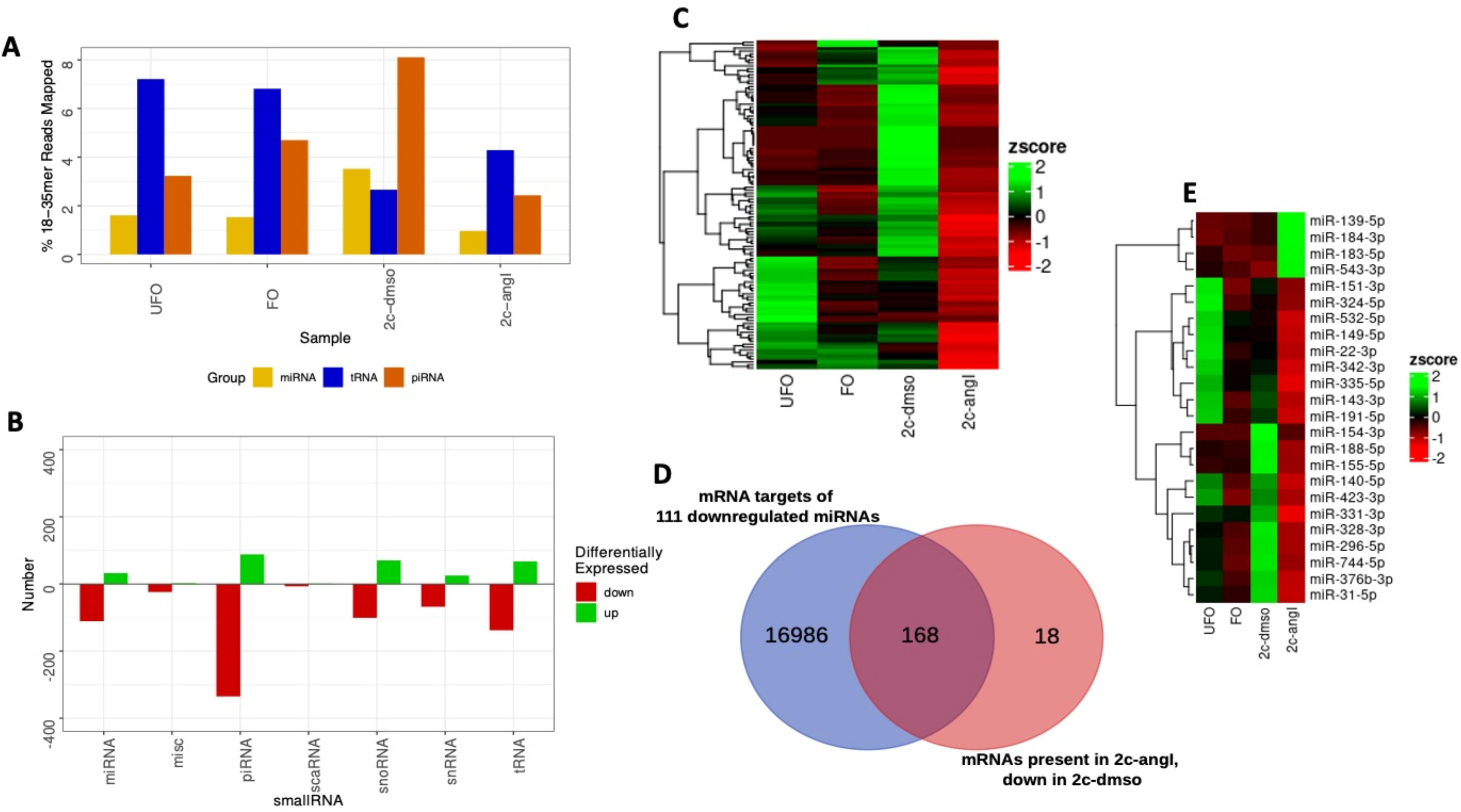
A) The bar graph shows the percentage of reads mapped to the genome for miRNAs, tsRNA, and piRNAs in UFO, FO, 2c-DMSO, and 2c-ANGi-treated conditions. B) The bar plot represents the differential small RNA expression pattern of miRNAs, misc-RNAs, piRNAs, scaRNAs, snoRNAs, snRNAs, and tsRNAsin 2c-ANGi treated m-embs compared to 2c-DMSO-treated m-embs in 24 hours. C) Heatmap depicts the upregulated miRNAs in 2c-DMSO compared to 2c-ANGi condition at 24 hrs, and further correlated this expression pattern with UFO and FO. Normalised read counts were used to generate this heatmap. D) Total miRNA predicted sites were detected for the miRNAs downregulated in the 2c-ANGi condition at 24hrs using the miRNA target prediction database (miRDB) for the mRNAs upregulated in 2c-ANGi/FO compared to 2c-DMSO/FO condition at 24hrs. E) Heatmap represents the miRNAs downregulated in 2c-ANGi condition at 24hrs and have miRNA predicted site for 168 mRNAs upregulated in 2c-ANGi/FO compared to 2c-DMSO/FO condition at 24hrs

We next performed differential expression analysis across the major small RNA classes, including miRNAs, tRNA-derived RNAs, piRNAs, snoRNAs, snRNAs, miscellaneous RNAs (misc_RNAs), and scaRNAs. Several small RNA classes showed reduced abundance in 2c-ANGi embryos compared with 2c-DMSO embryos at 24 h (Figure 5B). Together, these observations indicate that inhibition of ANG is associated with broad alterations in the small RNA landscape during early embryonic development. While these data do not distinguish between direct effects of ANG on small RNA processing and secondary effects resulting from developmental arrest, they raise the possibility that ANG activity contributes to the establishment or maintenance of specific small RNA populations during the 2-cell stage.

### Angiogenin inhibition alters miRNA expression associated with maternal RNA turnover

We next examined miRNAs that were differentially expressed in 2c-DMSO relative to 2c-ANGi embryos at 24 h and compared their expression patterns across UFO, FO, and 2c-DMSO stages (Figure 5C, Supp Table S3B). Many of these miRNAs showed reduced abundance in FO compared with UFO, followed by increased abundance in 2c-DMSO embryos. These dynamics suggest that distinct miRNA populations undergo stage-specific changes following fertilization and during the transition to the 2-cell stage. Such changes could reflect degradation of maternally deposited miRNAs followed by accumulation of newly expressed miRNAs; however, the present data do not distinguish newly transcribed miRNAs from maternally or paternally contributed species.

To examine the potential relationship between ANG-sensitive miRNAs and mRNA turnover, we predicted targets of 111 miRNAs that were downregulated in 2c-ANGi relative to 2c-DMSO embryos using miRDB. This analysis identified 16,986 predicted target sites. We then compared these predicted targets with the 186 transcripts retained or enriched in 2c-ANGi embryos described in Figure 3B. Of these 186 transcripts, 168 were predicted targets of one or more of the ANG-sensitive miRNAs (Figure 5D). Although target prediction alone does not establish direct miRNA-mediated regulation, the substantial overlap suggests a potential relationship between altered miRNA abundance and the persistence of transcripts following ANG inhibition. Several of these miRNAs were predicted to target multiple transcripts within the retained mRNA set (Figure 5E, Supp Table S3C), raising the possibility that coordinated changes in miRNA populations may contribute to mRNA turnover during early embryonic development.

Based on their expression dynamics across UFO, FO, 2c-DMSO, and 2c-ANGi samples, these miRNAs could be broadly classified into three groups (Figure 5E). miR-group 1, comprising miR-151-3p, miR-324-5p, miR-532-5p, miR-149-5p, miR-22-3p, miR-342-3p, miR-335-5p, miR-143-3p, and miR-191-5p, showed relatively high abundance in UFO, reduced levels following fertilization, and modest increases in 2c-DMSO embryos. This pattern is consistent with dynamic regulation of these miRNAs during the maternal-to-zygotic transition. miR-group 2, comprising miR-154-3p, miR-188-5p, and miR-155-5p, showed relatively similar abundance in UFO and FO but increased abundance in 2c-DMSO embryos. Their persistence following fertilization followed by increased abundance at the 2-cell stage suggests that these miRNAs may participate in post-transcriptional regulation during the period associated with zygotic genome activation. The last group, miR-group 3, including miR-140-5p, miR-423-3p, miR-331-3p, miR-328-3p, miR-296-5p, miR-744-5p, miR-376b-3p, and miR-31-3p, showed higher abundance in UFO, reduced abundance in FO, and subsequent enrichment in 2c-DMSO embryos. This biphasic pattern suggests distinct regulation before and after fertilization and is consistent with a potential role for these miRNAs during the 2-cell stage.

We further classified the 168 predicted mRNA targets according to the DBTMEE database. These transcripts were distributed across several developmental expression categories, with prominent representation of maternal RNA, MGA, minor ZGA, maternal-to-MGA, major ZGA, minor-ZGA-to-MGA/4-cell transient, 2-cell transient, maternal-to-ZGA, and 1-cell transient groups (Supp Table S3C, Supp Table S2A). Collectively, these analyses reveal stage-specific changes in miRNA abundance that are altered following ANG inhibition and identify a substantial predicted overlap between ANG-sensitive miRNAs and transcripts associated with the maternal-to-zygotic transition. These findings are consistent with a potential contribution of miRNA-mediated post-transcriptional regulation to maternal RNA turnover, although direct regulation of these transcripts by the identified miRNAs remains to be experimentally established.

### Angiogenin inhibition is associated with altered tsRNA and piRNA profiles during early embryogenesis

We next examined whether ANG inhibition was associated with changes in tsRNA abundance during early embryonic development. Heatmap analysis was performed for 18–35 nt tsRNAs differentially represented between 2c-ANGi and 2c-DMSO embryos at 24 h, and their abundance was compared with FO and UFO stages (Supp Figure 4A). A subset of tsRNAs abundant in UFO showed reduced levels in 2c-DMSO embryos, whereas several tsRNAs enriched in FO were either retained or reduced at the 2-cell stage. These patterns indicate extensive remodelling of the tsRNA population following fertilization.

A distinct subset of tsRNAs showed increased abundance in 2c-DMSO embryos compared with FO and UFO but reduced abundance following ANG inhibition. Thus, ANG inhibition disrupts the normal stage-associated dynamics of specific tsRNA populations. Whether these changes reflect direct ANG-dependent processing of tRNAs or secondary consequences of altered embryonic progression remains to be determined.

To further characterize ANG-sensitive tsRNAs, we examined the top 20 tsRNAs that were increased or decreased in 2c-ANGi relative to 2c-DMSO embryos at 24 h (Supp Figure 4B). tsRNAs enriched following ANG inhibition were derived predominantly from glycine, cysteine, alanine, serine, leucine, and pseudogene-encoded tRNAs. In contrast, tsRNAs derived from several other tRNA species, including phenylalanine, aspartate, leucine, proline, serine, threonine, isoleucine, arginine, tryptophan, glycine, glutamine, methionine, valine, tyrosine, and pseudogene tRNAs, were reduced following ANG inhibition (Supp Figure 4B). These findings demonstrate that ANG inhibition is associated with selective rather than uniform changes in tsRNA populations during the 2-cell stage.

We also examined the piRNA population. Previous studies in hamster embryos have shown that disruption of PIWI proteins can impair early embryonic development, supporting an important role for the piRNA pathway during preimplantation development [23]. We therefore analyzed piRNAs that were reduced in 2c-ANGi relative to 2c-DMSO embryos at 24 h and compared their abundance across UFO and FO stages (Supp Figure 4C). Many of these piRNAs showed increased abundance in 2c-DMSO embryos relative to UFO and FO but failed to show a comparable increase following ANG inhibition. This observation indicates that the normal accumulation of a subset of piRNAs at the 2-cell stage is altered when ANG activity is inhibited.

Analysis of the top 20 differentially represented piRNAs further identified substantial changes between 2c-ANGi and 2c-DMSO embryos (Supp Figure 4D). The abundance of these piRNAs also showed associations with maternal mRNA levels and with changes in stress-granule markers, tsRNAs, and miRNAs following ANG inhibition. These relationships suggest coordinated remodelling of multiple RNA regulatory pathways during the developmental arrest induced by ANG inhibition, although their functional interdependence remains to be established.

Overall, our findings demonstrate that ANG inhibition is accompanied by extensive changes in miRNA, tsRNA, and piRNA populations during the 2-cell stage. Together with the observed defects in maternal RNA clearance, these data support a model in which ANG activity is associated with the normal reorganization of post-transcriptional regulatory networks during the maternal-to-zygotic transition. Further experiments will be required to determine which small RNA changes represent direct consequences of ANG activity and which arise secondarily from developmental arrest.

### RNA granule and RISC-associated markers in mouse embryos following angiogenin inhibition

To investigate whether altered maternal RNA degradation (MRD) following angiogenin inhibition is associated with changes in RNA granule-associated pathways, we examined the expression of markers associated with stress granules, P-bodies, and the RISC complex in embryos treated with ANGi for 24 h and 47 ± 1 h. Several RNA granule- and RNA turnover-associated factors, including Tudor, TIAR, TIAL, Dcp1a, and Ago2, showed altered expression patterns between control and ANGi-treated embryos (Figure 4C, 4D; Supp Figure 3A, 3B), suggesting that ANG inhibition is accompanied by changes in components of the post-transcriptional RNA regulatory machinery.

We next examined the spatial association of the RNA granule-associated protein Tudor with the maternal transcripts Reep4 and Esrrb following ANGi treatment for 24 h (Supp Figure 5A, 5B). Qualitative analysis revealed increased colocalization of Tudor with Reep4 and Esrrb transcripts in FO and 2c-ANGi embryos compared with 2c-DMSO embryos at 24 h. These observations suggest that the persistence of maternal transcripts following ANG inhibition is associated with their increased localization to Tudor-positive RNA granules. One possibility is that localization within these RNA granules influences the accessibility of maternal transcripts to RNA turnover pathways during early embryonic development. However, whether Tudor-positive granules directly regulate transcript stability or degradation, and how this process relates to the altered small RNA profiles observed following ANG inhibition, remains to be established.

## Discussion

The maternal-to-zygotic transition (MZT) represents a critical developmental transition during which control of embryonic development shifts from maternally deposited RNAs and proteins to products of the zygotic genome. This transition involves extensive maternal RNA degradation (MRD) together with activation of zygotic transcription. Maternal RNA clearance is generally mediated through two overlapping mechanisms: maternal-factor-mediated decay (M-decay), which can occur independently of zygotic transcription, and zygotic genome activation (ZGA)-dependent decay (Z-decay) [1,3,20]. Despite considerable progress in defining these pathways, how maternal RNA turnover is coordinated with other post-transcriptional regulatory mechanisms during early mammalian development remains incompletely understood.

Our findings identify angiogenin (ANG) activity as an important component associated with this transition. Inhibition of ANG resulted in developmental arrest predominantly at the 2-cell stage and was accompanied by impaired clearance of a subset of maternal transcripts. These changes were further associated with altered expression of small RNAs and RNA granule-associated factors. Together, these observations suggest that ANG activity contributes to the coordinated RNA remodeling that accompanies progression through the 2-cell stage.

The developmental phenotype observed following ANG inhibition is particularly relevant in the context of ZGA. Abe et al. [3] demonstrated that inhibition of minor ZGA from the mid-1-cell to early 2-cell stage results in developmental arrest of more than 90% of mouse embryos at the 2-cell stage, whereas inhibition during the period associated with major ZGA produces a comparatively lower frequency of 2-cell arrest. These findings established that minor ZGA is important for subsequent major ZGA and developmental progression. Minor ZGA has also been implicated in chromatin remodeling and epigenetic changes that prepare the embryo for major ZGA [3]. Our observation that ANG inhibition results in 2-cell arrest together with persistence of maternal transcripts is consistent with disruption of processes associated with the MZT. However, whether ANG directly regulates ZGA or primarily influences RNA turnover during this transition remains to be determined.

Maternal RNA degradation involves multiple RNA-binding proteins and post-transcriptional regulatory complexes. Factors including Ago2, Dcp1a, SMAUG, BTG4, YAP1/TEAD4, and PABPN1L have been implicated in recognition, stabilization, or degradation of maternal transcripts during early embryogenesis [1,2,24–29]. We observed altered expression of several RNA granule- and RNA turnover-associated factors, including Tudor, TIAR, TIAL, Dcp1a, and Ago2, following ANG inhibition. Importantly, we also observed increased association of the maternal transcripts Reep4 and Esrrb with Tudor-positive RNA granules in ANGi-treated embryos compared with 2c-DMSO controls. These observations raise the possibility that ANG-dependent RNA remodeling during the MZT involves changes in the organization or dynamics of messenger ribonucleoprotein (mRNP) complexes.

The functional relationship between mRNP granules and RNA degradation is likely to be dynamic. P-bodies, stress granules, and related mRNP assemblies share components and can exchange RNAs and proteins, with their assembly state influencing RNA localization, translation, storage, and degradation [30–34]. Importantly, the presence or abundance of microscopically detectable RNA granules does not necessarily correlate directly with RNA degradation. Several studies have suggested that larger cytoplasmic P-body assemblies can function as sites of RNA storage, whereas RNA decay may involve smaller or more dynamic mRNP complexes [33–36]. In this context, the increased association of Reep4 and Esrrb transcripts with Tudor-positive granules following ANG inhibition is intriguing. One possibility is that altered mRNP organization affects the accessibility of maternal transcripts to RNA turnover machinery, thereby contributing to their persistence. Direct measurements of RNA stability and granule dynamics will be required to test this model.

Another notable observation from our study is the extensive remodeling of the small RNA landscape during the 2-cell stage and its disruption following ANG inhibition. miRNAs, endogenous siRNAs, piRNAs, and tsRNAs have been implicated in post-transcriptional regulation during development, although their relative contributions to mammalian MZT remain incompletely defined. Small RNA-mediated mechanisms have been linked to maternal RNA turnover in several organisms, including Drosophila, zebrafish, Xenopus, and mammals [1,29], suggesting that small RNAs represent an evolutionarily recurrent component of post-transcriptional regulation during early development.

Our analysis identified a population of miRNAs whose abundance was reduced following ANG inhibition. Target prediction revealed that 168 of the 186 transcripts retained or enriched following ANG inhibition contained predicted target sites for one or more of these miRNAs. Although computational target prediction does not establish direct regulation, this substantial overlap raises the possibility that altered miRNA abundance contributes to the persistence of a subset of maternal and developmentally regulated transcripts. The stage-specific expression patterns of these miRNAs further suggest that distinct miRNA populations may participate at different phases of the maternal-to-zygotic transition.

ANG inhibition was also associated with substantial changes in tsRNA and piRNA populations. Given the known ribonuclease activity of ANG and its established ability to generate tRNA-derived fragments under specific cellular contexts, the altered tsRNA profiles are particularly noteworthy. However, the present experiments do not establish whether the observed tsRNAs are generated directly through ANG-mediated cleavage in early embryos. Similarly, the reduced abundance of a subset of piRNAs following ANG inhibition suggests a relationship between ANG activity and the piRNA landscape but does not establish direct regulation of piRNA biogenesis. These changes may therefore reflect direct RNA processing activities of ANG, indirect effects on RNA regulatory pathways, or consequences of developmental arrest at the 2-cell stage.

Taken together, our findings support a model in which ANG activity is associated with coordinated remodeling of multiple RNA regulatory processes during the maternal-to-zygotic transition (Figure 6). We propose that ANG-dependent changes in small RNA populations and mRNP organization may influence the accessibility and turnover of maternal and other developmentally regulated transcripts, thereby facilitating progression beyond the 2-cell stage. Inhibition of ANG disrupts this RNA regulatory environment and is accompanied by persistence of maternal transcripts and developmental arrest. An important next step will be to determine which of these effects result directly from ANG-mediated RNA processing and which arise indirectly from altered developmental progression.

In summary, this study identifies ANG as a previously underappreciated component of RNA regulation during early mouse embryogenesis and establishes a connection between ANG activity, maternal RNA clearance, small RNA dynamics, and RNA granule organization. Rather than acting through a single RNA pathway, our findings suggest that ANG may contribute to the broader reorganization of the post-transcriptional landscape that accompanies MZT. Defining the direct RNA substrates of ANG and determining how ANG activity interfaces with small RNA pathways and mRNP dynamics will be important for establishing the molecular mechanisms through which ANG contributes to early embryonic development.

## Methodology

### Mouse embryo collection protocol

C57BL/6NJ mice were used for the study. B6NJ_2020 & B6NJ_2022 females 24-29 days old were superovulated using PMSG (5IU) and after 48 hours using HCG (5IU). Following HCG injection, female mice were introduced to male mice either immediately or after 5 hrs [37], and fertilised m-embryos were collected after 15-16 hours of HCG injection. Briefly, female mice were sacrificed using cervical dislocation and the oviduct was collected and further dissected to collect the cumulus cells. Next, cumulus cells were digested using hyaluronidase enzyme in M2 media to release the m-embs. Mouse embryos were washed in M2 media 4-5 times and then cultured in KSOM media. M2 media and KSOM media were prepared according to the Cold Spring Harbour Protocols with slight modifications. To study the influence of ANG on the progression of m-embs, They were treated with ANGi for 24hrs and 47±1 hrs and snap frozen for RNA isolation and fixed in 4% PFA for immunofluorescence study. ANGi N65828 was used in our study as described by Kao et al. (2002) [11]. As ANGi was dissolved in DMSO, and therefore equal amount of DMSO was added to the control condition.

### In-situ and Immunofluorescence study of mouse embryos

For the in situ and immunofluorescence (IF) study, m-embs were collected and fixed in 4% PFA. Further, m-embs were fixed on coverslip-embedded dishes using poly-d-lysine coating and subsequently stored in either PFA or used immediately. For in-situ hybridization, RNA probes were prepared using in-vitro transcription reaction by mixing 1ug HindIII digested double stranded DNA with 2ul 10X transcription buffer (Thermo Scientific), 1X FITC (Roche), T7 RNA polymerase (Thermo Scientific), 5mM DTT, RNAse inhibitor (1U/ul, Thermo Scientific), 1ul Inorganic pyrophosphatase and 10mM MgCl2 in 20ul reaction incubated at 37°C for 4 hrs. Further, RNA probes were further purified by adding 3M sodium acetate pH 5.4 and 95% ethanol with overnight incubation at -80oC and then centrifuged at 14000rpm, 4°C for 40 min and washed with 70% ethanol for 15 min.

For combined IF and in-situs experiments, m-embs were stained for IF first and then used for in-situ hybridization using 500bp FITC and DIG-labelled RNA probes. Briefly, m-embs were permeabilized using 0.5% triton-x-100 for 1hr followed by 2hr incubation in blocking buffer composed of 5% BSA and 0.1% PBST (PBS + 0.1% tween-20). Further, m-embs were incubated in primary antibody prepared in blocking buffer for 3-4 hours. Then, m-embs were washed with PBS 2-3 times and a secondary antibody with Hoechst 3342 was added for 1-2 hr. After incubation, m-embs were washed with PBST 2-3 times, and coverslips were either mounted on glass slides or proceeded for in-situ hybridization. For in-situ hybridization, m-embs were first incubated in a hybridization buffer (50% formamide, 2X SSC, 5% dextran sulphate, 2.5% Denhadht solution for 45 min at 45oC in a hybridization chamber. RNA probes were prepared in a hybridization buffer containing 70% formamide and incubated at 70°C for 15 min followed by 2 min incubation on ice. After 45 mins, m-embs were incubated with a hybridization buffer containing RNA probes for 3hr at 45°C in a hybridization chamber. After hybridization, m-embs were washed 2-3 times with 2X SSC and 1.5X SSC to remove the non-specific binding and coverslips were mounted on glass slides using mounting media for FITC-RNA probes. For DIG-labelled probes, m-embs were washed 2-3 times with 2X SSC after hybridization and then followed by incubation with anti-DIG POD antibody for 1-2 hr at room temperature (RT) in 10mM Tris, pH 7.5 and 2X SSC buffer. After incubation, m-embs were washed twice with 2X SSC buffer and incubated with secondary cy5 cy3 or FITC tyramide in 0.003% H_2_O_2_ for 1 hour at RT. After incubation, m-embs were washed 2-3 times with 2X SSC and 1.5X SSC and mounted on glass slides using mounting media.

### RNA isolation from mouse embryos

50-80 m-embs were collected from 4-5 breeding pairs of animals in the presence and absence of ANGi at 24 hours, and 47±1 hr. 150ul of trizol was added to the m-embs and pipetted several times. Further, RNA was extracted from the aqueous layer by adding 1/5 volume of chloroform and centrifuged at 14000 rpm for 40 mins. Next, isopropanol was added to the aqueous layer and centrifuged at 14000 rpm for 40 min to pellet the RNA. RNA pellet was washed twice with 70% ethanol, air dried and resuspended in 15ul of nuclease-free water (NFW). RNA was quantified using nanodrop, and Qubit reading and RNA integrity was analysed on bioanalyzer.

### Small RNA sequencing library preparation

Small RNA sequencing libraries were prepared using the SMARTer smRNA-seq kit for Illumina according to the manufacturer’s instructions. For small RNA library preparation, 1ng of total RNA was collected from 4-5 different sets of experiments (n=26 mice) as described by Ko et al. (2023) [38] for pooled sequencing. Briefly, an artificial poly-A tail was added in polyadenylation reaction using 0.25 ul of Poly(A) Polymerase (2 U/μl) at 16 °C for 5 min and further subjected to cDNA synthesis using oligo dT primers and PrimeScript RT (200 U/μl) in a thermocycler at 42 °C for 60 min, 70 °C for 10 min and 4 °C hold. Following cDNA synthesis, full-length Illumina adapters were added to the cDNA by PCR using 2ul of SeqAmp DNA Polymerase with forward and reverse primers in a thermocycler at 98 °C-1 min, and 27 cycles at 98 °C -10s, 60 °C-5s, 68 °C-10s and hold at 4 °C. Further, the PCR product was purified by a PCR clean-up kit, and the libraries were double size selected using the AMPure XP beads-based method according to the above small RNA library preparation kit and quantified and analysed using a bioanalyzer.

### mRNA sequencing library preparation

mRNA sequencing libraries were prepared using NEBNext® Single Cell/Low Input RNA Library Prep Kit for Illumina according to the manufacturer’s instructions. 20pg of total RNA was taken in duplicate from 4-5 different set of experiments (n=2; 13 mice each set) as described by Ko et al. (2023) [38] for pooled sequencing and reverse transcribed in a thermocycler using NEBNext Single Cell RT Enzyme Mix and NEBNext Template Switching Oligo for 90 min at 42°C and the reaction was terminated by incubating the mixture at 70°C for 10min. cDNA was amplified using NEBNext Single Cell cDNA PCR Master Mix and NEBNext Single Cell cDNA PCR Primer at 98°C for 45s for 1-cycle and further extended using 21 cycles at 98°C for 10s, 62oC for 15s, 72°C for 3 min and finally extended at 65°C for 5min and kept at hold at 4°C. The amplified cDNA was cleaned up using Ampure XP beads. The cDNA was further used for fragmentation and adapter ligation using NEBnext ultra II FS enzyme mix and NEBnext ultra II ligation mix, respectively. Adapter-ligated DNA was further cleaned up by using Ampure XP beads, and PCR enrichment was performed for the same. After PCR enrichment, the libraries were size selected using the Ampure XP beads-based method and quantified on a bioanalyzer.

### RNA-Seq data analysis

RNA from UFO, 1c, 2cand 4c stages (n=1) were sequenced on an Illumina Hiseq 2500 instrument in a paired-read, 100-base mode. FastQC v0.11.5 was used to perform the initial quality check and the adapter sequence was then trimmed from the demultiplexed reads using cutadapt version 2.10. The trimmed reads were next aligned to the mouse genome (GRCm39) using STAR v2.5.0a using default parameters. The Ensembl GRCm39 annotation was used to quantify transcripts using featureCounts v1.5.0-p1. Read count normalisation and differential expression analysis was done using the R package ‘NOISeq’ (norm=“tmm”, pnr=0.2, nss=5, v=0.02). Only those genes that were differentially expressed with a probability score greater than 0.9 were considered for analysis.

mRNA seq libraries from the ANGi experiment (n=2, pooled sample) were sequenced using the Illumina NovaSeq6000 instrument in a paired-read, 51-base mode. Read trimming, mapping and counting were performed using cutadapt, STAR aligner and featureCounts, as before. Read count normalisation and differential expression analysis was done using the R package ‘DESeq2’. Only those genes that showed at least a two-fold change with q<0.05 in at least one of the comparisons (dmso-2c_vs_FO or angI-2c_vs_FO) were considered for further analysis. The classification for the genes expressed during mouse early embryogenesis was obtained from the DBTMEE database (version 2.0). Gene set enrichment analysis was performed using the GSEA webserver. All the plots were generated using R packages.

### Small RNA-Seq data analysis

Small RNA libraries were sequenced using the Illumina NovaSeq6000 instrument in a single-read, 51-base mode.

Read trimming was performed using cutadapt and reads that were 18 to 35 bases long were selected for further analysis. The trimmed reads were next aligned to the mouse genome (GRCm39) using STAR v2.5.0a using default parameters. The Gencode (version M32) annotation set was used to quantify the small RNAs (tRNA, misc_RNA, rRNA, scaRNA, snoRNA, snRNA). For piRNA, annotation from the RNAcentral database (version 22.0) and for miRNA, the annotation from mirBase (version 22.1) was used for quantification using featureCounts. Read count normalisation and differential expression analysis was done using the R package ‘NOISeq’ as before. Only those genes that had a probability score greater than 0.9 were considered for further analysis. The miRNA targets were obtained from the TargetScan database (release 8.0; *Conserved_Family_Info.txt*). All the plots were generated using R packages.

### Statistical analysis

Statistical analysis for the in-situ hybridisation and immunofluorescence experiments was performed by the two-sample t-test method by calculating mean and standard deviation as described by Xu et al. (2017) [39]. The p-values were calculated by using MedCalc online software ( MedCalc Software, 2024) [40]. P-values are represented with asterisk < 0.05 (*), < 0.01 (**), < 0.001 (***), < 0.0001 (****).

## Author contributions

Conceptualization, D.P and T.M.; Methodology, S.S, N.H, D.P, A.J, A.S.K; PKV Software, N.H, S.S.; Validation, S.S, N.H, D.P.; Formal Analysis, S.S, N.H, D.P.; Investigation, S.S, N.H, D.P.; Resources, D.P.; Data Curation, N.H, S.S.; Writing – Original Draft Preparation, S.S.; Writing – Review & Editing, S.S, N.H, D.P.; Visualization, D.P.; Supervision, D.P.; Project Administration, D.P.; Funding Acquisition, D.P and T.M. All authors have read and agreed to the published version of the manuscript.

## Institutional Review Board statement

The animal work was conducted according to the Institutional Animal Committee (Animal Care and Resource Center at National Center for Biological Sciences) guidelines and approved by the Ethics Committee.

## Informed consent statement

Non applicable

## Data availability statement

The raw data for mRNA and small RNA sequencing is submitted to NIH Bioproject database with the accession number ’PRJNA1189325. The BioProject and associated SRA metadata are available at https://dataview.ncbi.nlm.nih.gov/object/PRJNA1189325?reviewer=11l2s0akffa7u59i0c7fiqa7r4 in read-only format.

---

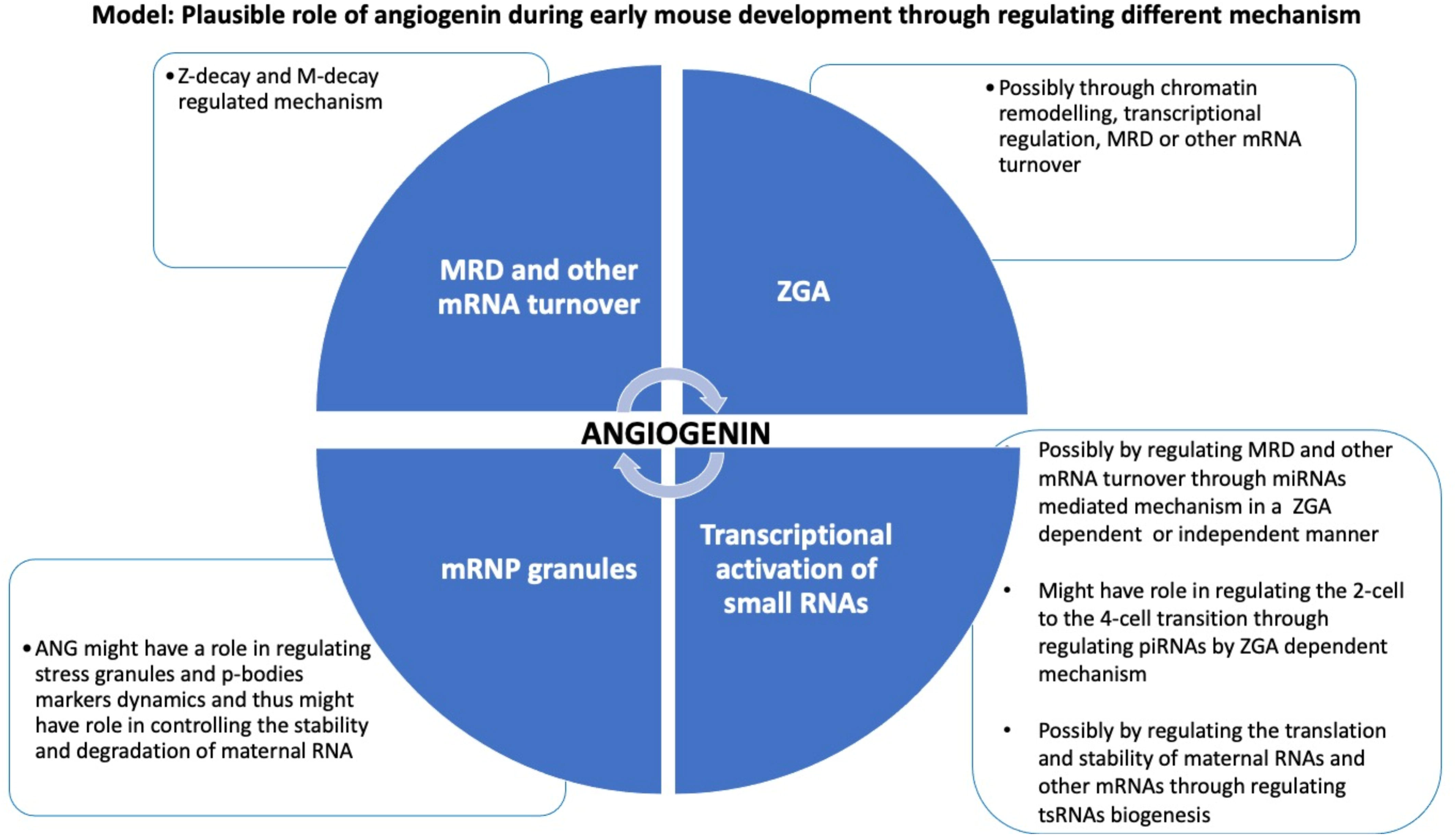

