## Supplementary Figures for "Angiogenin activity regulates RNA remodeling during the maternal-to-zygotic transition in early mouse embryogenesis"

**Supp Figure 1**

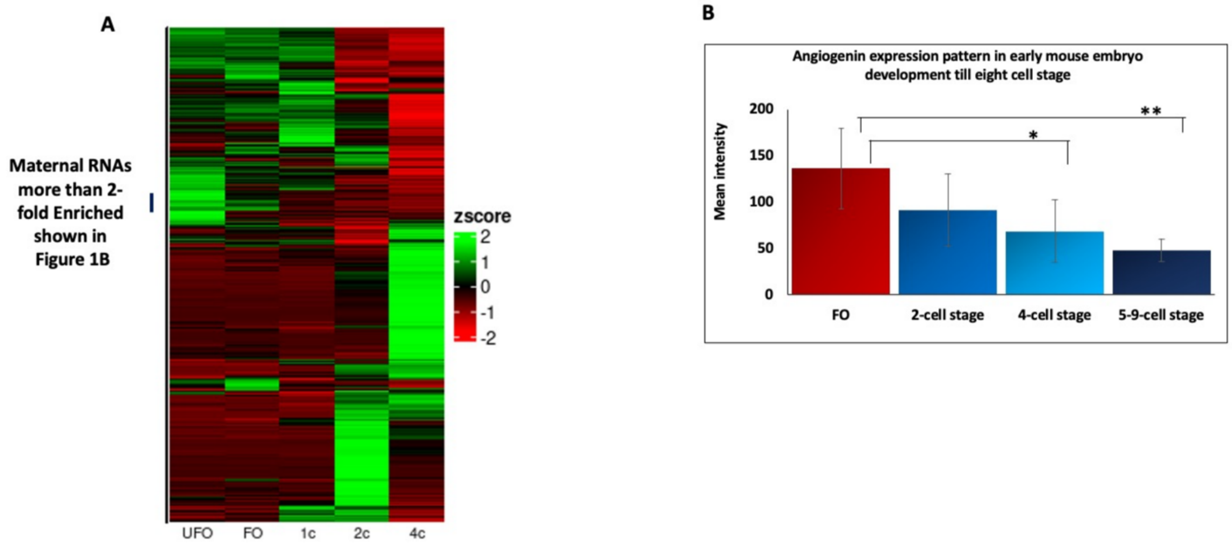


**Supp Figure 1:**

A) Heatmap representation of transcriptomic analysis during early developmental stages of m-embs, such as unfertilized oocytes (UFO), fertilized oocytes (FO), late 1-cell stage (1c), 2-cell stage (2c), and 4-cell stage (4c) m-embs.

B) The bar graph denotes the mean intensity of ANG expression in FO, 2-cell stage, 4-cell stage, and 8-cell stage of m-embs calculated from the confocal images using ImageJ software.

**Note: UFO = unfertilized oocytes, FO = fertilized oocytes, L-1c = late 1-cell, 2c = 2-cell stage, and 4c = 4-cell stage**

*p-value are represented with asterisk <0.05 (*), <0.01 (**), <0.001 (***), <0.0001 (****).*

**Supp Figure 2**

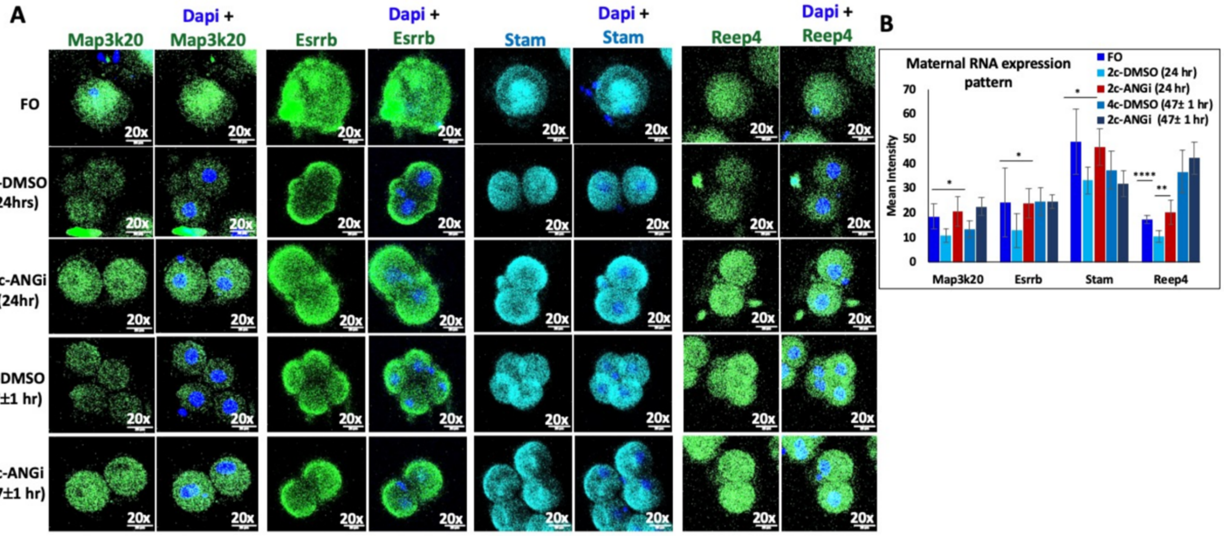


**Supp Figure 2:**

A) To affirm RNA-seq data, In-situ hybridization was done for mRNAs, including Map3k20, Esrrb, Stam, and Reep4, upregulated in 2c-ANGI/FO compared to 2c-DMSO/FO condition. The m-embs were cultured in the presence and absence of ANGi for 24 hours, and 47±1 hrs compared to DMSO control condition, and in situ-hybridization fluorescent intensity was detected using confocal microscopy shown by representative images. The experiment was done on the animals (n=15) obtained from the pooled samples of 3-4 breeding pair mice and approx. 15-20 mouse embryos were analysed for each condition.

B) The bar graph represents the mean intensity of Map3k20, Esrrb, Stam, and Reep4 mRNA expression in FO, 2c-DMSO, 4c-DMSO, and 2c-ANGi at 24hrs and 47±1 hrs. The mean intensity was calculated from the confocal images (n=4-8) using ImageJ software.

*p-value are represented with asterisk <0.05 (*), <0.01 (**), <0.001 (***), <0.0001 (****).*

### Supp Figure 3

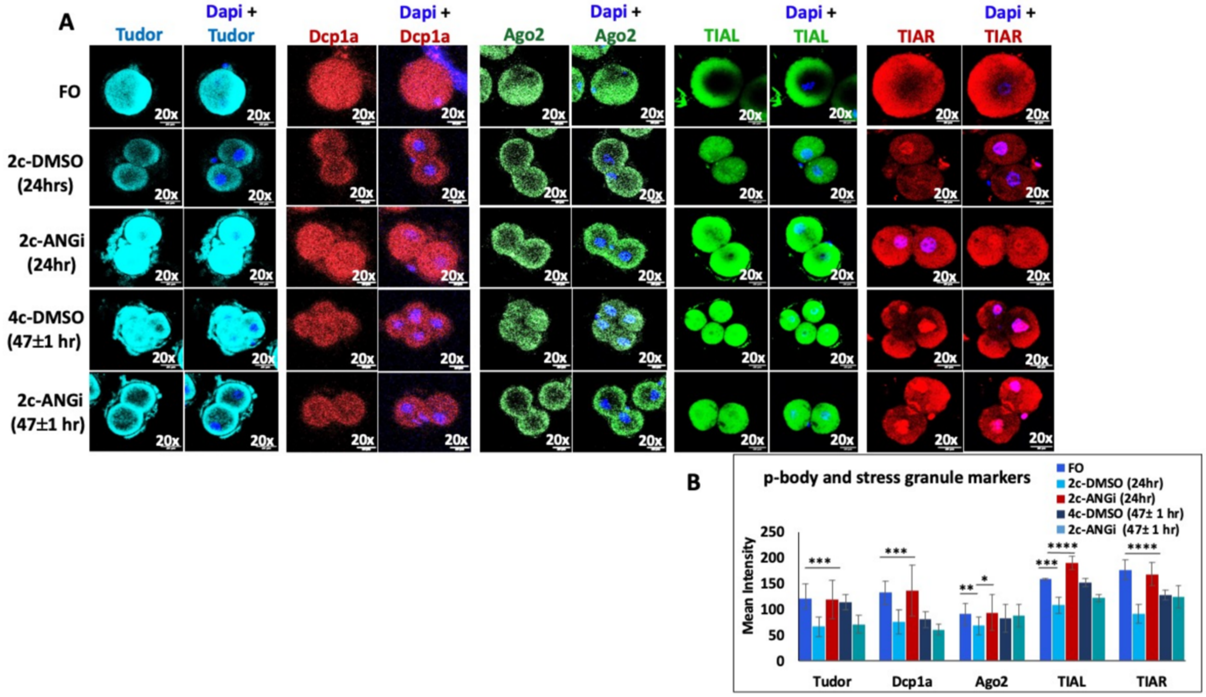


**Supp Figure 3:**

A) Immunofluorescence detection of stress granule markers (Tudor, TIAL and TIAR), p-body marker (Dcp1a), and RISC complex marker (Ago2) in FO, 2c-DMSO, 4c-DMSO, and 2c-ANGi m-embs at 24hrs and 47±1 hrs shown by representative images.

B) Bar plot shows the mean intensity of Tudor, Dcp1a, Ago2, TIAL and TIAR expression in FO, 2c-DMSO, 4c-DMSO, and 2c-ANGi treated m-embs at 24hrs and 47±1 hrs. The mean intensity was calculated from the confocal images (n=8-16) using ImageJ software. The experiment was repeated n=3 times.

*p-value are represented with asterisk <0.05 (*), <0.01 (**), <0.001 (***), <0.0001 (****).*

**Supp Figure 4**

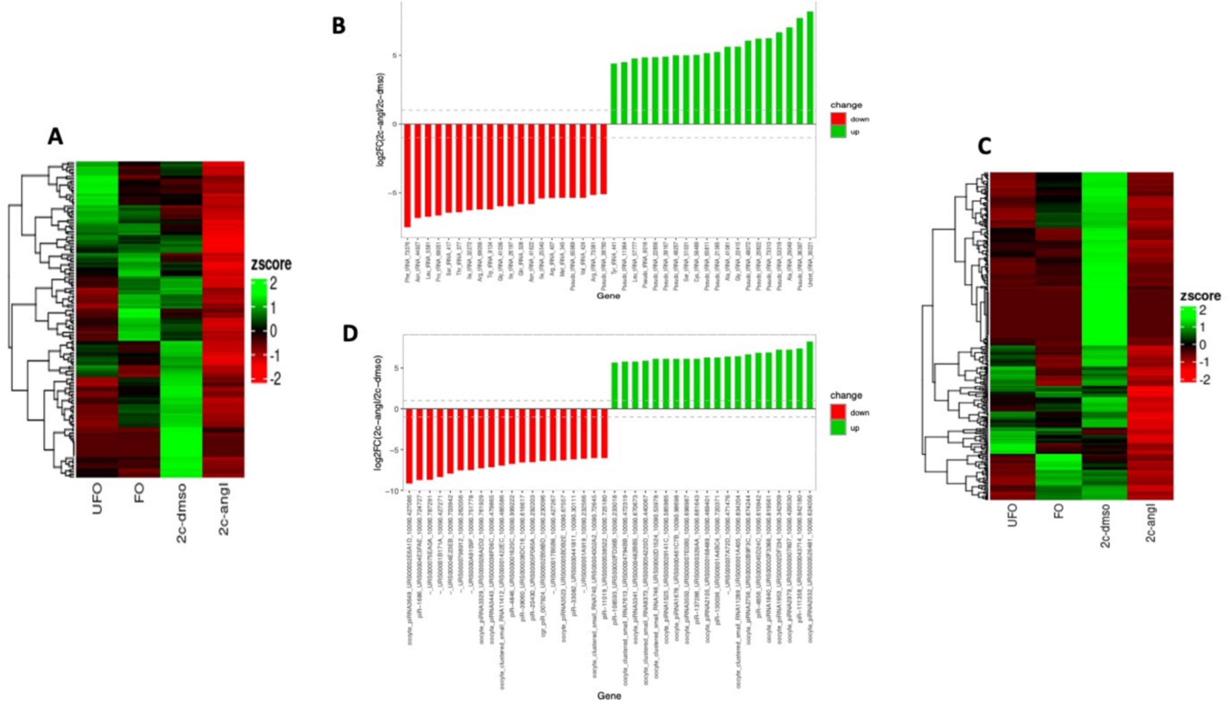


**Supp Figure 4:**

A) Heat map represents the 18-35 mer tsRNAs upregulated in 2c-DMSO condition compared to 2c-ANGi condition and further compared with UFO and FO.

B) Bar plot represents the top 20 tsRNA downregulated/upregulated in 2c-ANGi condition compared to 2c-DMSO condition at 24hr.

C) Heat map represents the 18-35 mer piRNAs upregulated in 2c-DMSO condition as compared to 2c-ANGi condition and further compared with UFO and FO.

D) Bar plot represents the top 20 piRNAs downregulated/upregulated in 2c-ANGi condition compared to 2c-DMSO condition at (24hr).

**Supp Figure 5**

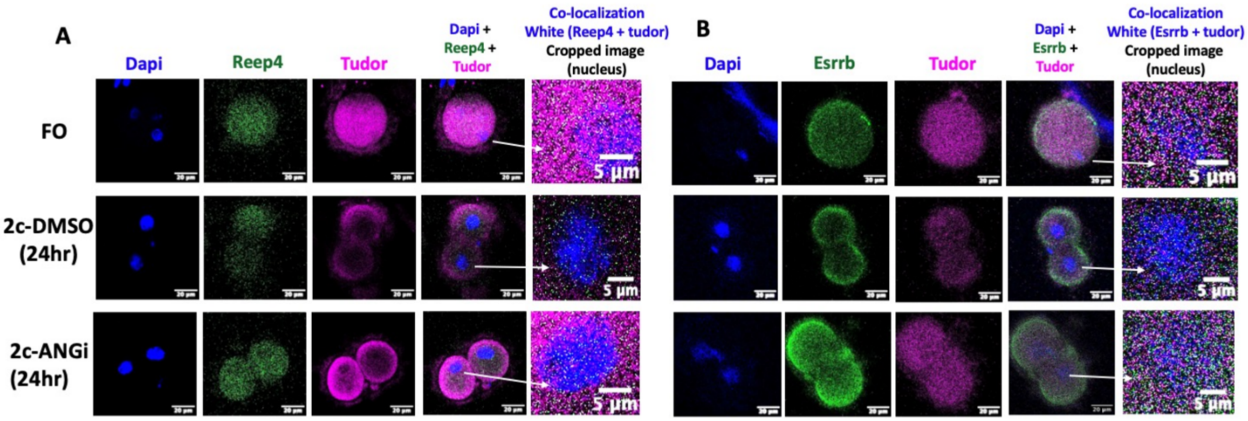


**Supp Figure 5:**

A) Co-localization pattern of Reep4, and Tudor in FO, 2c-DMSO, and 2c-ANGi at 24hrs

B) Co-localization pattern of Esrrb and Tudor in FO, 2c-DMSO, and 2c-ANGi at 24hrs.
